# Histone serves as an eat-me signal to induce RAGE-mediated phagocytosis

**DOI:** 10.1101/2024.09.03.610921

**Authors:** Yuqing Li, Xiaoman Zhou, Yan Yang, Congcong Du, Yi-shi Liu, Zijie Li, Hideki Nakanishi

## Abstract

The receptor for advanced glycation end products (RAGE) is a multiligand receptor that can induce phagocytosis in both professional and nonprofessional phagocytes. We found that histones are another ligand for RAGE. Binding between histones and RAGE is increased when DNA is attached to histones. While histones are chromosomal proteins in healthy cells, they are exposed to the cell surface as a complex with DNA when cells undergo apoptosis. The phagocytosis of apoptotic cells by either professional or nonprofessional phagocytes is enhanced when histones are present on the surface of apoptotic cells. Thus, histones serve as eat-me signals. In *RAGE* knockout cells, the phagocytosis of apoptotic cells was not influenced by the removal of histones, indicating that RAGE is required for the removal of histones from histone-presenting cells. In RAGE knockout mice, wound healing and removal of apoptotic cells from wound sites are delayed, suggesting that RAGE-mediated phagocytosis functions under physiological conditions.

## Introduction

Phagocytosis is an endocytic process in which relatively large particles greater than 0.5 µm in size are internalized by cells via a receptor-mediated mechanism. Phagocytosis is mediated primarily by professional phagocytes including macrophages, dendritic cells, and neutrophils [1]. Since professional phagocytes present various phagocytic receptors, including those for antibodies and complements, they can internalize a variety of particles and macromolecules [2]. In addition to professional phagocytes, many other types of cells, including epithelial cells and fibroblasts, can perform phagocytosis; these cells are termed nonprofessional phagocytes [3]. Compared with those of professional phagocytes, the number of phagocytic receptors present in nonprofessional phagocytes is limited. However, they can internalize certain particles, such as apoptotic cells [3–5].

Apoptotic cells are rapidly phagocytosed so that they are removed before the release of proinflammatory cellular components [6]. Apoptotic cells present unique phagocytic ligands, called eat-me signals, that can activate phagocytic receptors [7]. For example, the exposure of phosphatidylserine (PS) to the outer leaflet of the plasma membrane in apoptotic cells is known to serve as an eat-me signal [8, 9]. PS is recognized by multiple phagocytic receptors either directly or indirectly [10]. Direct PS-recognizing receptors include T-cell immunoglobulin and mucin domain (TIM) family receptors [11], brain angiogenesis inhibitor 1 (BAI1) [12], stabilin-1 [13], stabilin-2 [14], and receptor for advanced glycation end-products (RAGE) [15].

RAGE is a member of the immunoglobulin superfamily proteins and was originally identified as an advanced glycation end products (AGE)-recognizing receptor that induces the proinflammatory signaling pathway [16, 17]. This receptor is known to bind multiple ligands; high mobility group box 1 (HMGB1), polynucleotides, and S100 proteins are reported as its ligands in addition to PS and AGE. Prolonged activation of RAGE causes chronic inflammation [18], which is involved in the progression of various disorders [19–21]. In addition to its ability to induce proinflammatory signals, RAGE can induce phagocytosis in both professional and nonprofessional phagocytes [15]. In professional phagocytes, RAGE has been shown to induce the clearance of apoptotic cells by binding to PS [15, 22].

In a previous study, we reported that particles bound to histones are targeted for RAGE-mediated phagocytosis [22], suggesting that histones are another ligand for RAGE. Histones are components of chromatin but they can be released from cells by cell lysis or by the formation of neutrophil extracellular traps (NETs) [23]. Extracellular histones exhibit harmful effects: first, they can damage the plasma membrane by direct binding; second, they serve as damage-associated molecular patterns (DAMPs) and activate proinflammatory signaling pathways [24]. TLR2, TLR4, and TLR9 are reported to be receptors that recognize extracellular histones [25, 26]. Intriguingly, several reports have shown that histones are also present on the surface of apoptotic cells [27, 28]. This observation may be contradictory since apoptosis is believed to be a form of death that does not induce inflammation. However, cell surface histones or nucleosomes may enhance the recognition and clearance of apoptotic cells. A previous study reported that cell surface histones are targeted by an opsonin ApoJ, which is known to induce phagocytosis by binding to LDL receptor-related protein (LRP) [29, 30]. In the present study, we demonstrated that cell surface histones are direct targets for RAGE-mediated phagocytosis. Histone-induced and RAGE-mediated phagocytosis is involved in the clearance of apoptotic cells under physiological conditions, such as in the wound repair process.

## Results

### Histones are ligands of RAGE

To verify that histones are ligands of RAGE, binding between the purified histone complex and RAGE was assayed in vitro. For this experiment, the extracellular part of RAGE termed RAGE^23-341^ (Fig 1A) was produced by HEK293 cells. The core histone complex purified from bovines was coprecipitated with RAGE^23-341^ (Fig. 1B). RAGE^124-341^ is an extracellular region of RAGE lacking the V domain (Fig. 1A). Compared with that of RAGE^124-341^, the number of histone complexes bound to RAGE^23-341^ was increased 2.4-fold (Fig. 1B). These results demonstrate that the histone complex directly binds to RAGE via the V domain. A previous study showed that histone-induced phagocytosis was enhanced when DNA was bound to histones [22]. Thus, we examined whether the binding between RAGE and the histone complex was improved when DNA was bound to histones. The DNA-bound histone complex, hereafter referred to as the DNA-histone complex, was prepared by incubation of a single-strand DNA (22 nucleotides) [22] molecule with the histone complex. As shown in Fig. 1C, the amount of the histone complex that precipitated with RAGE^23-341^ increased when the DNA was bound. We also examined the binding between RAGE^23-341^ and recombinant human histone H3.1. Similar to the purified histone complex, recombinant histone H3.1 was coprecipitated with RAGE^23-342^, and the binding of these proteins decreased when the V domain was removed (Fig. S1A). The binding of histone H3.1 to RAGE^23-341^ was improved when DNA was bound to histone H3.1 (Fig. S1B).

**Figure 1.**
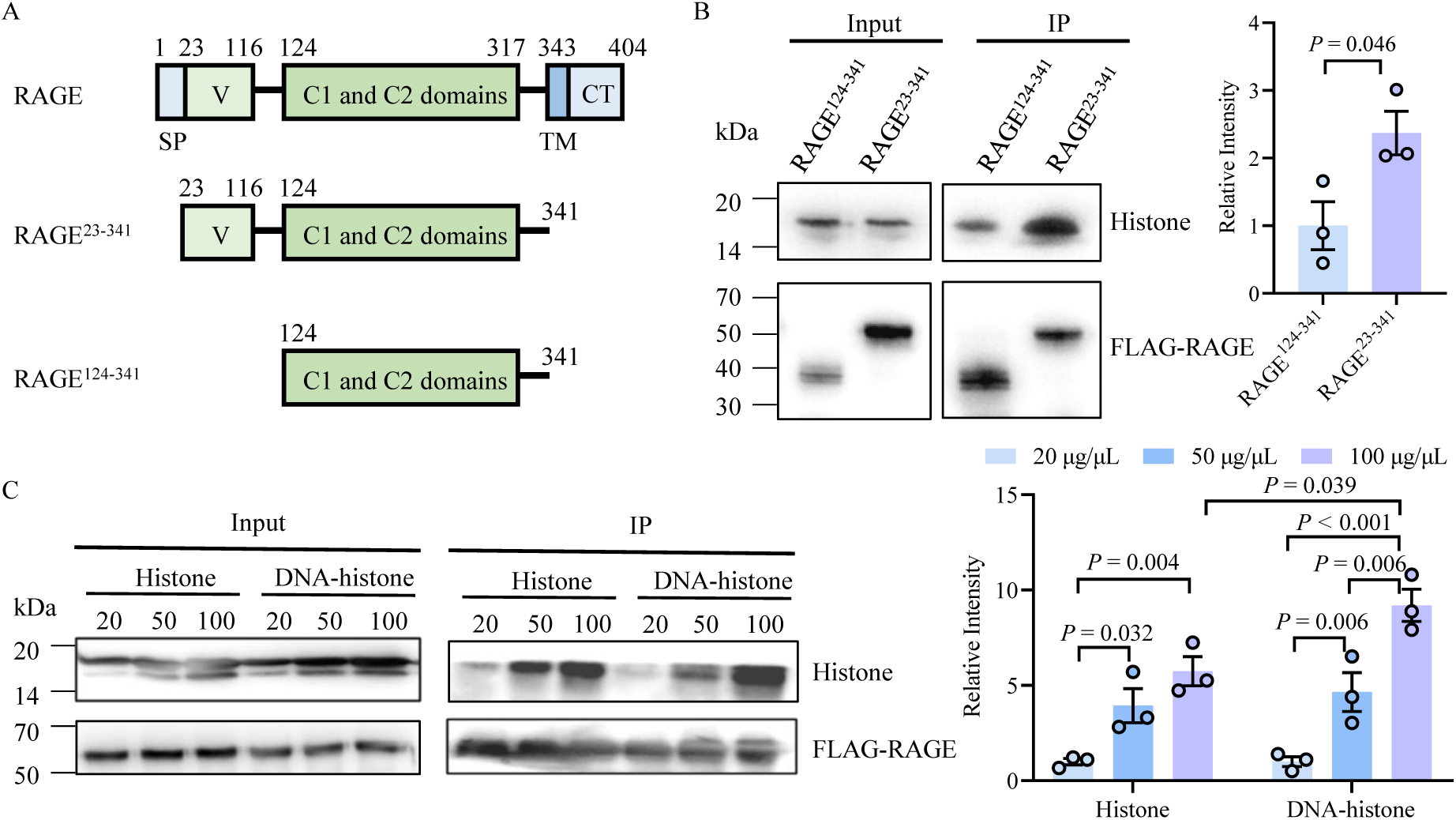
Histones bind to RAGE. (A) Schematic diagrams of wild-type and truncated RAGE proteins. SP, signal peptide; V, V domain; TM, transmembrane domain; CT, cytosolic tail. (B) The histone complex was precipitated with RAGE^23-341^-FLAG or RAGE^124-341^-FLAG attached to agarose beads. Left panels: Inputs and precipitated proteins were detected with an anti-FLAG antibody. Right panel: Quantification of histones precipitated with truncated RAGEs. Histones precipitated with RAGE^124-341^-FLAG were defined as 1, and the relative intensity of histones precipitated with RAGE^23-341^-FLAG is shown. (C) Left panels: RAGE^23-341^-FLAG was incubated with histones or DNA-histones at the indicated concentrations. Inputs and precipitated proteins were detected with an anti-FLAG antibody. Right panel: Relative intensities of histones precipitated with RAGE^23-341^-FLAG. The intensity of the precipitated histones when the assay was performed at a histone concentration of 20 μg/μl was defined as 1. Data were presented as mean ± SEM. n = 3. *P* values were derived from an unpaired two-tailed Student’s t-test.

### Histones are attached to the apoptotic cell surface via DNA

Previous studies have shown that nucleosomes are exposed on the surface of apoptotic cells [28]. Accordingly, histones were detected by immunofluorescence microscopy in unpermeabilized apoptotic Jurkat cells (Fig. S2). Apoptotic Jurkat cells were prepared via staurosporine treatment, and the induction of apoptosis was verified via the detection of annexin V and caspase-3 (Fig S3). Flow cytometry analysis also revealed that the degree of histone staining was increased by the induction of apoptosis (Fig. 2A). Histone staining in apoptotic cells was decreased by proteinase treatment (Fig. 2B). Furthermore, histones were released by DNase treatment (Fig. 2C and D). Notably, the purified histone complex could bind to proteinase-treated apoptotic cells (Fig. 2B), presumably because proteinase-treated apoptotic cells remain cell surface DNA which can accommodate the histone complex. In line with this notion, histones did not bind to apoptotic cells treated with either proteinase or DNase (Fig. 2E). Recombinant histone H3 also bound to proteinase-treated apoptotic cells (Fig. 2F).

**Figure 2.**
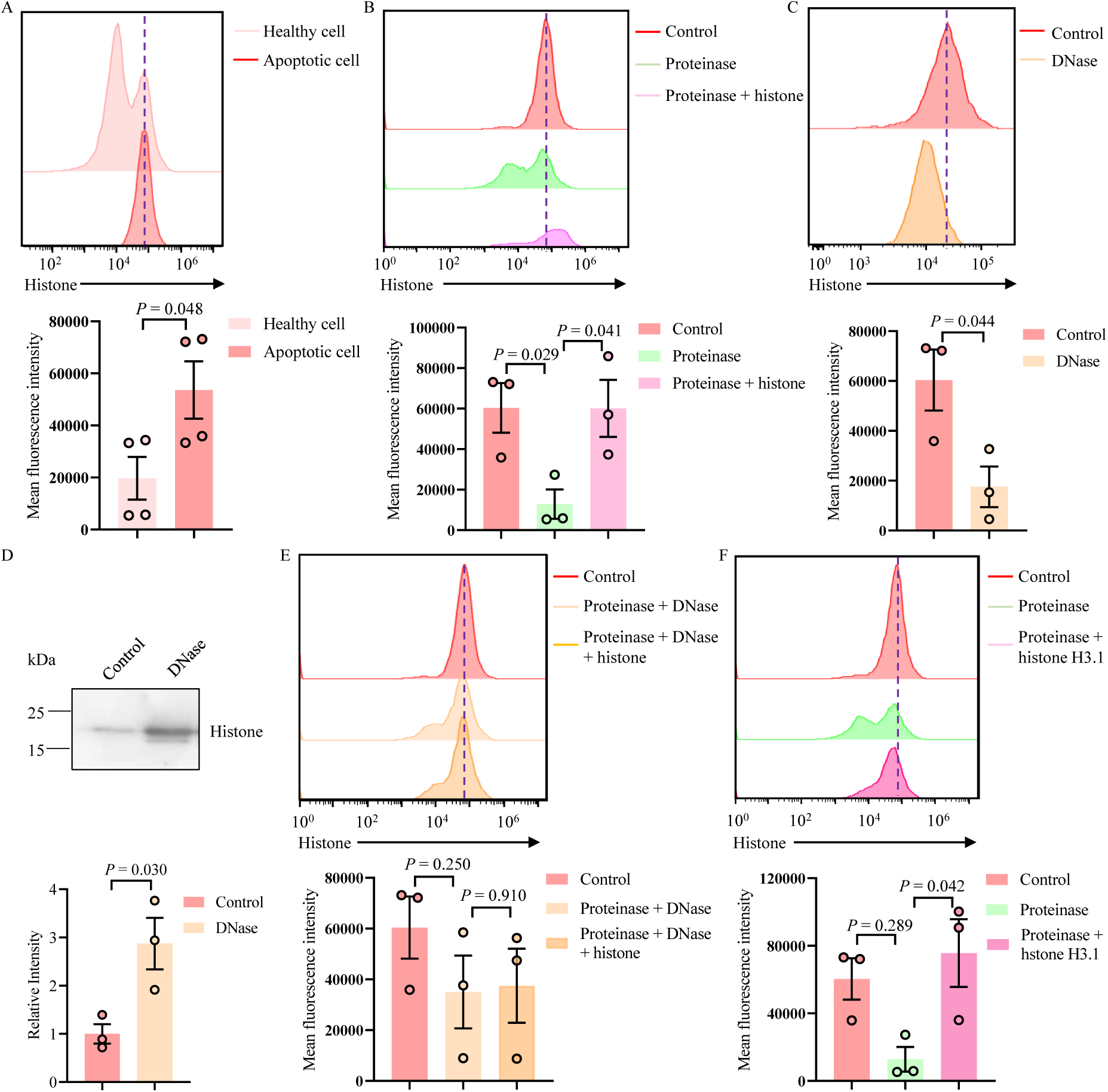
DNA-histones on the surface of apoptotic cells. (A) Cell surface histones were labelled with an anti-histone H3 antibody in healthy and apoptotic Jurkat cells. These cells were analyzed by flow cytometry. The mean fluorescence intensity values are shown in the bottom panel. (B) Apoptotic Jurkat cells were treated with or without proteinase. Proteinase-treated cells were incubated with or without the purified histone complex. Cell surface histones were analyzed by flow cytometry. The mean fluorescence intensity values are shown in the bottom panel. (C) Cell surface histones present on apoptotic cells treated with or without DNase were analyzed by flow cytometry. The mean fluorescence intensity values are shown in the bottom panel. (D) Apoptotic Jurkat cells were treated with or without DNase, and histones released into the supernatant were detected by western blotting. Quantification of released histones is shown in the bottom. The intensity of histone released from apoptotic Jurkat cells without DNase treatment was defined as 1. (E) Apoptotic cells were treated with or without proteinase and DNase. The cells treated with proteinase and DNase were incubated with or without the purified histone complex. The surface histones present on these cells were analyzed via flow cytometry. The mean fluorescence intensity values are shown in the bottom panel. (F) Apoptotic cells were treated with or without proteinase. Proteinase-treated cells were incubated with or without recombinant histone H3.1. The surface histones present on these cells were analyzed via flow cytometry. The mean fluorescence intensity values are shown in the bottom panel. Data are presented as mean ± SEM. n = 4 (A); n = 3 (B to F). *P* values were derived from an unpaired two-tailed Student’s t-test.

### Histones serve as ligands to induce RAGE-mediated phagocytosis

Given that histones are ligands for RAGE, we hypothesized that histone molecules present on the cell surface of apoptotic cells could serve as an eat-me signals to induce RAGE-mediated phagocytosis. This hypothesis was first assessed in an epithelial cell line, HEK293T, because RAGE serves as a primary receptor to mediate phagocytosis in this cell line [22]. A phagocytosis assay was performed via a flow cytometry-based method; phagocytic cells were sorted to determine whether they harbored GFP-expressing Jurkat cells. As reported previously [22], HEK293T cells internalized apoptotic Jurkat cells (Fig. S4A). The fraction of HEK293T cells that internalized apoptotic Jurkat cells was decreased by 71% in *RAGE*-knockout (*RAGE*^-/-^) cells (Fig. 3A). Phagocytosis deficiency in *RAGE*^-/-^ HEK293T cells was reversed by the transfection of *RAGE* (Fig. 3A). To examine whether phagocytosis was competitively inhibited by soluble histone molecules, a phagocytosis assay was performed with HEK293T cells preincubated with the histone complex. As shown in Fig 3B, the percentage of internalized Jurkat cells decreased by 53% following pretreatment with the histone complex. The removal of cell surface histones by proteinase or DNase treatment caused a decrease in the number of apoptotic cells internalized by HEK293T cells (Fig. 3C and D). Importantly, the phagocytosis levels of proteinase-treated apoptotic cells were improved when the apoptotic cells were preincubated with the histone complex (Fig. 3C). However, the internalization of apoptotic cells treated with proteinase and DNase was not improved by preincubation with the histone complex (Fig. 3E), most likely because apoptotic cells treated with these enzymes lack the ability to accommodate histones (Fig. 2E). To confirm the results of these flow cytometry-based phagocytosis assays, the internalization of apoptotic cells was also quantified via microscopy assay, in which the number of HEK293T cells containing Jurkat cell-derived fragments (GFP fluorescence) in endocytic compartments was counted via fluorescence microscopy. Both FACS and microscopy assays revealed similar results for the internalization of apoptotic cells by wild-type and *RAGE*^-/-^ HEK293T cells: internalized apoptotic cells were decreased in *RAGE*^-/-^ HEK293T cells compared with wild-type cells, and the internalization levels of apoptotic cells were decreased by proteinase treatment, which was reversed by the addition of histones (Fig S4B). Thus, a flow cytometry-based quantification method was used for the remaining phagocytosis assays in this study. To further assess whether cell surface histones can induce phagocytosis, we used an anti-histone antibody to block the interaction between histones and receptors. As shown in Fig. 3F, the internalization of apoptotic Jurkat cells by HEK293T cells was inhibited when the apoptotic cells were preincubated with an anti-histone antibody. Notably, RAGE-KO cells remain the ability to phagocytose apoptotic cells, as the percentage of *RAGE*^-/-^ cells that internalized apoptotic cells decreased by 35% when the apoptotic cells were treated with proteinase (Fig. 3C). However, in *RAGE*^-/-^ cells, the internalization levels of proteinase-treated apoptotic cells were not increased by the binding of histones (Fig. 3C). In addition, the level of phagocytosis in *RAGE*^-/-^ cells was not altered by anti-histone antibody treatment (Fig. 3F). These results support the notion that histone-induced phagocytosis is mediated by RAGE in HEK293T cells.

**Figure 3.**
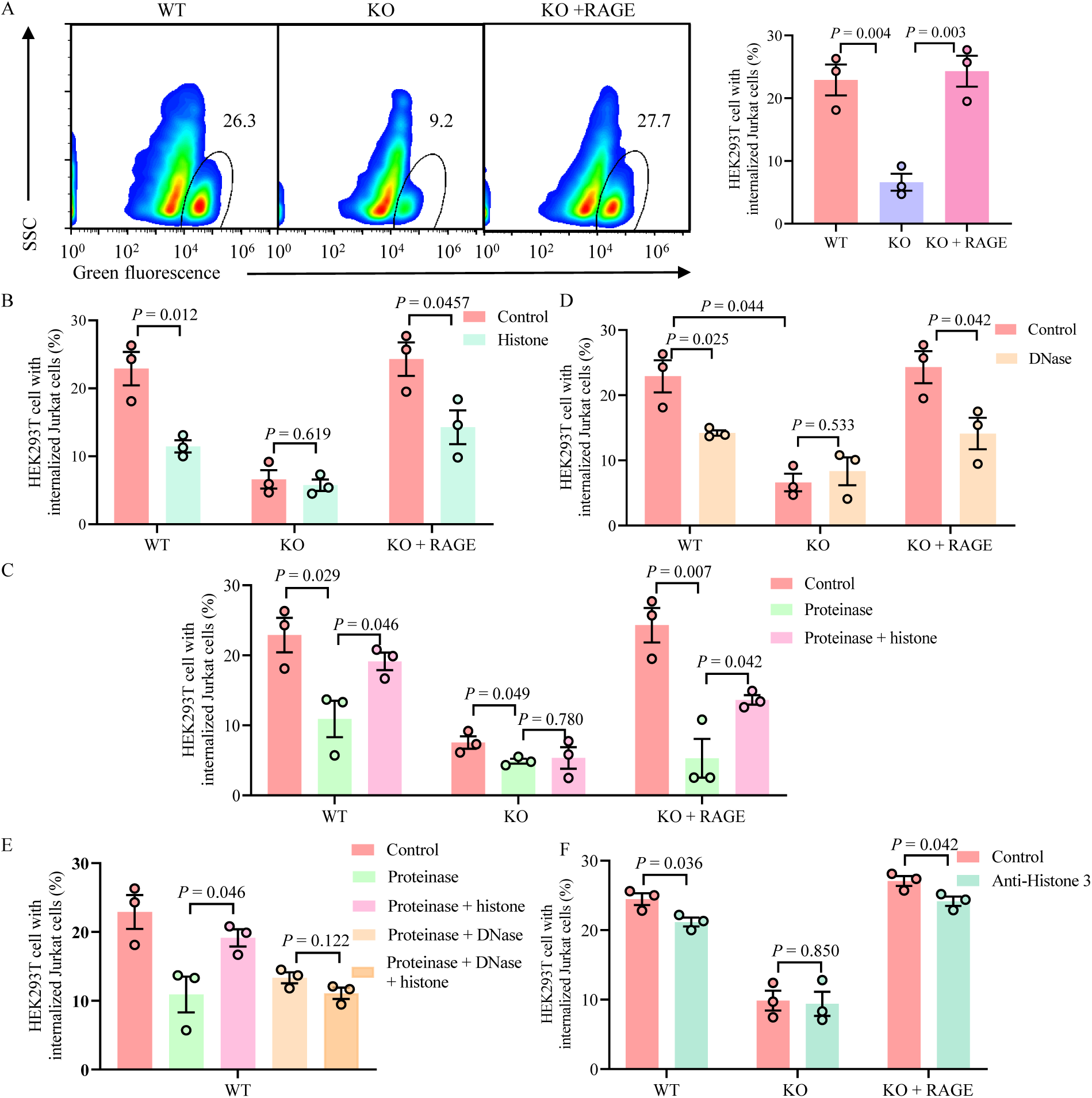
Cell surface histones on apoptotic cells induce RAGE-mediated phagocytosis in HEK293T cells. (A) Apoptotic Jurkat cells expressing GFP were incubated with wild-type HEK293T cells, *RAGE*^-/-^ cells, or *RAGE*^-/-^ cells transfected with *RAGE*. HEK293T cells that internalized Jurkat cells were analyzed via flow cytometry. The percentages of HEK293T cells harboring Jurkat cells are shown in the right panel. (B) Wild-type HEK293T cells, *RAGE*^-/-^ cells, or *RAGE*^-/-^ cells transfected with *RAGE* were preincubated with or without histones. These cells were subsequently incubated with apoptotic Jurkat cells, and the percentages of HEK293T cells harboring Jurkat cells were measured via flow cytometry. (D) Apoptotic cells were treated with or without DNase. These cells were incubated with wild-type HEK293T cells, *RAGE*^-/-^ cells, or *RAGE*^-/-^ cells transfected with *RAGE*, and the percentages of HEK293T cells harboring Jurkat cells were measured via flow cytometry. (E) Apoptotic cells were treated with proteinase or proteinase and DNase. Then, the samples were incubated with or without histones. These cells were incubated with wild-type HEK293T cells, and the percentages of HEK293T cells harboring Jurkat cells were measured via flow cytometry. (F) Apoptotic cells were pretreated with anti-histone H3 antibody or nonspecific control IgG. The cells were incubated with wild-type HEK293T cells, *RAGE*^-/-^ cells or *RAGE*^-/-^ cells transfected with *RAGE*. The percentages of HEK293T cells harboring Jurkat cells were measured via flow cytometry. Data were presented as mean ± SEM. n = 3. *P* values were derived from an unpaired two-tailed Student’s t-test.

For the above-described phagocytosis assays, HEK293T and apoptotic cells were incubated for 0.5 h. However, we found that the effects of DNase or proteinase were dependent on the incubation time of apoptotic cells and HEK293T cells. As shown in Fig. S5A, the effects of DNase became negligible when the assay was performed with a 1 h incubation. Compared with that of the 0.5 h incubation, the effect of proteinase treatment was also decreased when the cells were incubated for 1 h (Fig. S5B). These results suggest that apoptotic cells treated with hydrolytic enzymes can be internalized via a histone-independent phagocytosis process. Thus, histone-induced and RAGE-mediated phagocytosis is likely involved in the rapid elimination of apoptotic cells.

### Histones induce RAGE-mediated phagocytosis in primary cells

To assess whether histone-induced phagocytosis occurs in vivo, a phagocytosis assay was performed with primary cells. Since RAGE is highly expressed in lung tissues [31, 32], pulmonary epithelial cells were used for these experiments. Apoptotic cells were prepared from primary thymocytes; thymocytes were labelled with PKH67 and apoptosis was induced by dexamethasone (Fig. S6). The assay was performed with an incubation time of 0.5 h. In the epithelial cells, fractions that internalized apoptotic cells were decreased by proteinase, DNase, or anti-histone antibody treatments (Fig. 4A, B, and C). The level of protein internalized by proteinase-treated apoptotic cells improved when histones were bound (Fig 4A). In pulmonary epithelial cells derived from *Rage*^-/-^ mice (Fig S7), the fraction of internalized apoptotic cells was decreased compared to that of wild-type cells (Fig 4A). In the knockout cells, the level of internalized apoptotic cells was not altered by anti-histone antibody treatment (Fig 4C). Furthermore, the internalization of proteinase-treated apoptotic cells was not improved by the binding of histones (Fig 4A). We also performed a phagocytosis assay with primary macrophages obtained from peritoneal fluid (Fig 4D). Phagocytosis was decreased in macrophages when histones were removed from apoptotic cells by proteinase (Fig 4E) or DNase (Fig 4F) treatment. In *Rage*^-/-^ macrophages, the level of phagocytosis was not decreased by these treatments (Fig 4E and F). The internalization of proteinase-treated apoptotic cells by wild-type, but not *Rage*^-/-^ macrophages was improved by the binding of histones (Fig 4E). These results suggest that histone-induced and RAGE-mediated phagocytosis occurs in macrophages.

**Figure 4.**
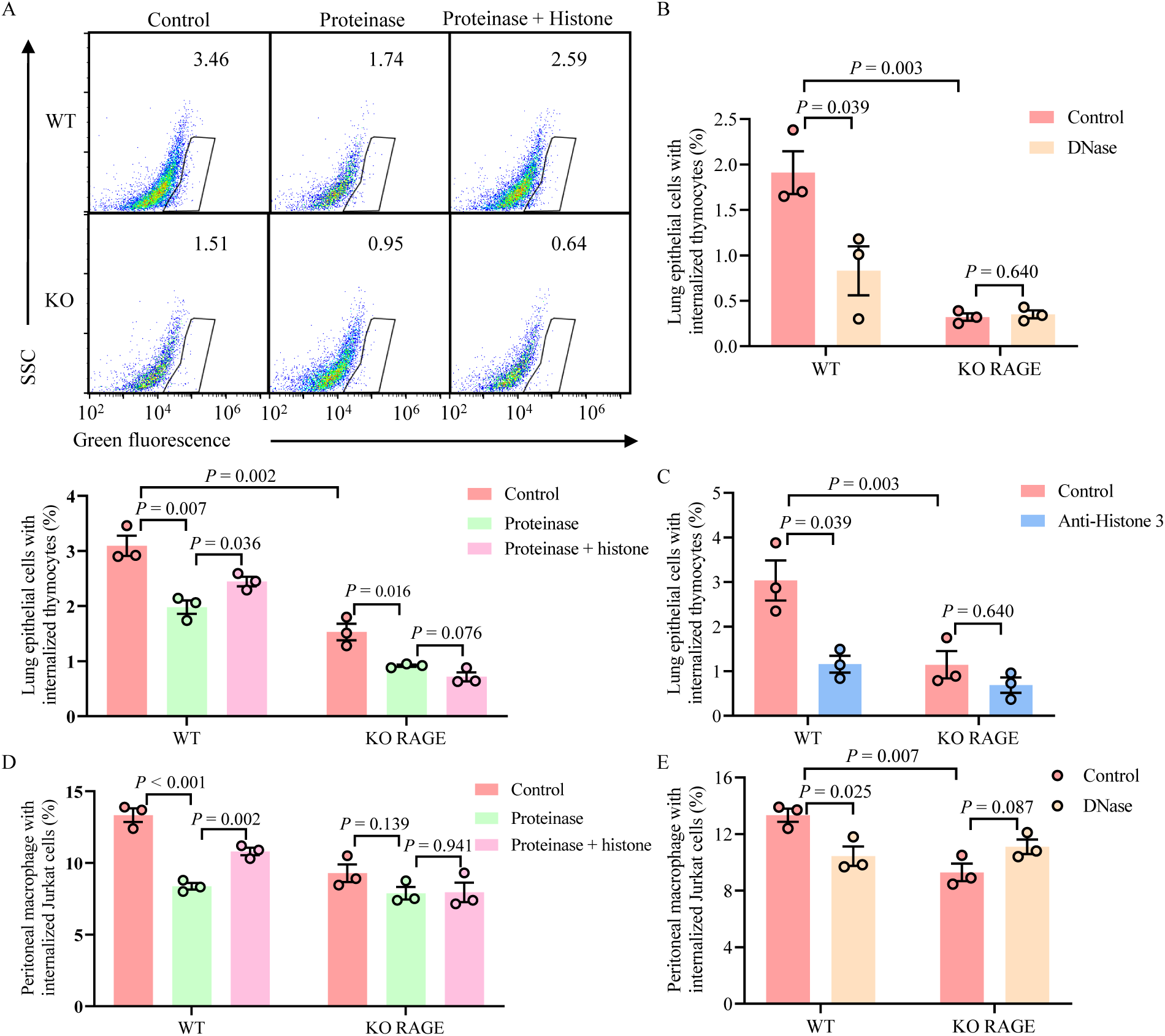
Histone-induced and RAGE-mediated efferocytosis occurs in mouse primary cells. (A) Apoptotic thymocytes labelled with PKH67 were treated with or without proteinase. Proteinase-treated cells were incubated with or without the purified histone complex. These cells were incubated with primary lung epithelial cells, and a phagocytosis assay was performed via flow cytometry. The percentages of lung epithelial cells harboring thymocytes are shown in the bottom panel. (B) Apoptotic thymocytes were treated with or without DNase. The internalization of these cells in lung epithelial cells was analyzed via flow cytometry. (C) Apoptotic thymocytes were pretreated with anti-histone H3 antibody or nonspecific control IgG. The internalization of these cells in lung epithelial cells was analyzed via flow cytometry. (D) Apoptotic thymocytes were treated with or without proteinase. Proteinase-treated cells were incubated with or without the purified histone complex. The internalization of these cells in peritoneal macrophages was analyzed via flow cytometry. (E) Apoptotic thymocytes were treated with or without DNase. The internalization of these cells in peritoneal macrophages was analyzed via flow cytometry. Data were presented as mean ± SEM. n = 3. *P* values were derived from an unpaired two-tailed Student’s t-test.

### RAGE is required to remove histone-bound cells during wound-healing

During the wound-healing process, the majority of neutrophils recruited to the wound site undergo apoptosis and are removed via phagocytosis [33]. To assess whether RAGE-mediated phagocytosis functions under physiological conditions, apoptotic cells present in wound sites were analyzed in wild-type and *Rage*^-/-^ mice. For this purpose, wounds were generated on the backs of the mice. Wound healing in *Rage*^-/-^ mice was slower than that in wild-type mice (Fig. 5A), which is consistent with a previous report [34]. Three days after wounding, the wound tissues were harvested, and the apoptotic cell fraction was analyzed via flow cytometry. In *Rage*^-/-^ mice, the number of apoptotic cell fractions (labelled with annexin V-PE) was greater in *Rage*^-/-^ mice than in wild-type mice (Fig. 5C). Neutrophils were sorted via CD45, CD11b and Ly-6G antibodies, and the number of apoptotic neutrophils (labelled with annexin V-PE) was also greater in *Rage*^-/-^ mice than in wild-type mice (Fig. 5D). Furthermore, the fractions of histone-presenting cells were greater in *Rage*^-/-^ mice than in wild-type mice (Fig 5E). These results indicate that RAGE-mediated phagocytosis is required for the removal of apoptotic cells from wound sites.

**Figure 5.**
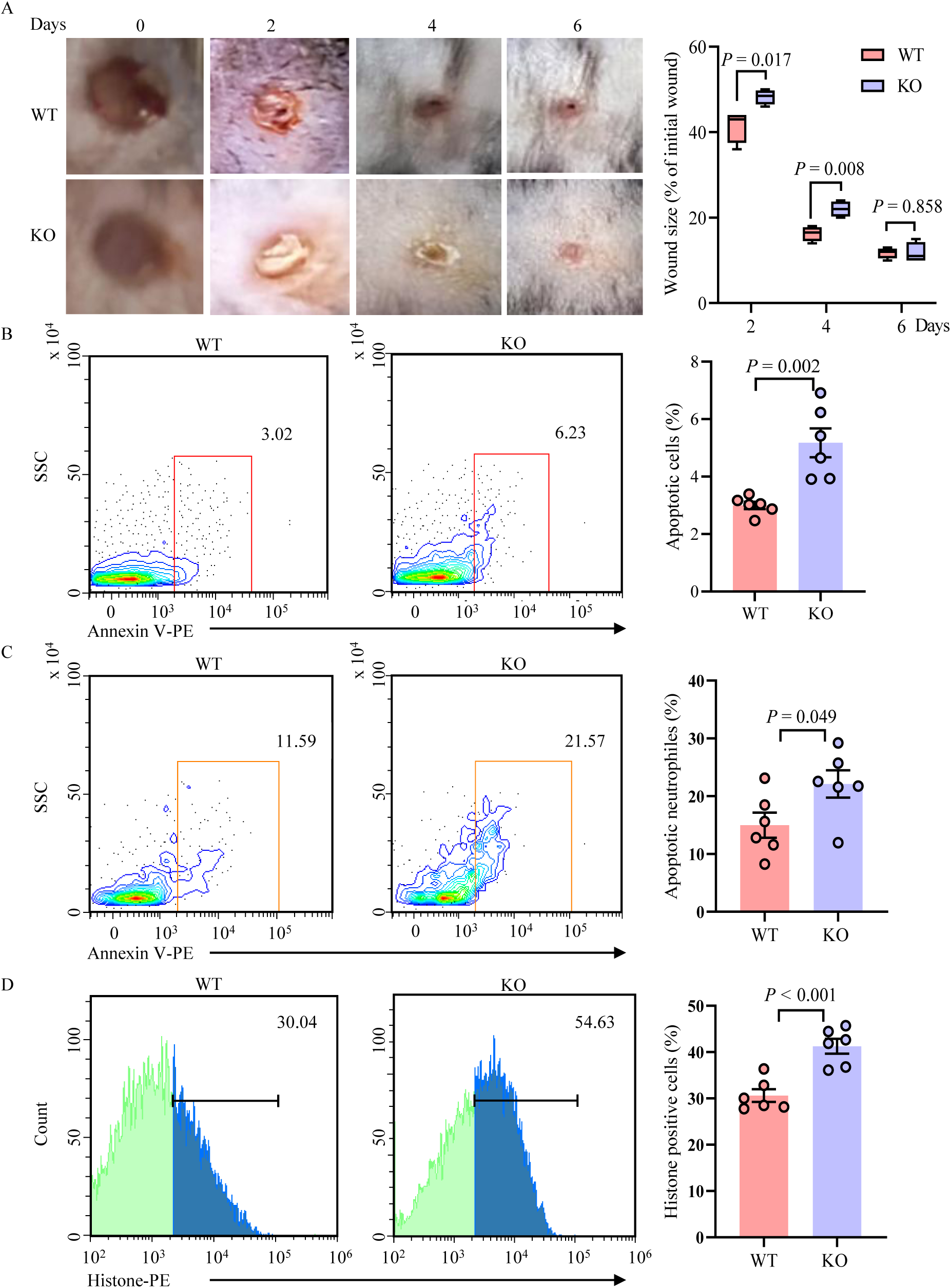
RAGE is involved in wound healing. Wounds were generated on the backs of the mice with a 3-mm punch. (A) Left panels: Representative images of wounds in wild-type and *Rage*^-/-^ mice. Right panel: Time course of the changes in the longest point spanning the length of the wound. In box plots, the center line denotes median, box edges encompass 25^th^ to 75^th^ percentiles, and whiskers span minimum to maximum values. n = 4. (B) Three days after wounding, wound tissues were collected, and cell suspensions were labelled with annexin V-PE to detect apoptotic cells. The cells were analyzed by flow cytometry. The percentages of apoptotic cells are shown in the right panel. (C) Neutrophils were sorted using CD45, CD11b and Ly-6G antibodies. Cell suspensions were labelled with annexin V-PE to detect apoptotic neutrophils. The cells were analyzed by flow cytometry. The percentages of apoptotic neutrophils are shown in the right panel. (D) Wound tissues were collected, and cell suspensions were labelled with an anti-histone H3 antibody. The cells were subsequently analyzed via flow cytometry. The percentages of labelled cells are shown in the right panel. Data were presented as mean ± SEM and n = 6 (B to D). *P* values were derived from an unpaired two-tailed Student’s t-test.

## Discussion

In the present study, we demonstrated that histones on apoptotic cells are directly recognized by RAGE in professional and nonprofessional phagocytes. Interference with their binding causes defects in the phagocytic removal of apoptotic cells, demonstrating that histones serve as eat-me signals to induce RAGE-mediated phagocytosis. Previous studies reported that histones induce LRP-mediated phagocytosis, whereas ApoJ is involved in this process as a histone-binding opsonin [29]. Thus, histones induce multiple phagocytosis pathways.

The mechanism by which histones are delivered to the cell surface during the apoptotic process remains elusive. Previous cytological analysis revealed that nucleosomes are exposed on the cell surface [28]. Consistent with this result, we found that histones are released from apoptotic cells under DNase treatment. Histone-induced and RAGE-mediated phagocytosis is increased when DNA is bound to histones [22]. In accordance with this synergetic effect between histones and DNA, the histone-DNA complex bound to RAGE with higher affinity than when bound to histones alone. Our results suggest that histones bind to the V domain in RAGE, via various ligands, including nucleotides [35–37]. Thus, the histone binding site in the V domain is distinct from the previously reported DNA-binding site, although further structural analyses are needed to clarify the mechanism by which RAGE can recognize multiple ligands.

Both professional and nonprofessional phagocytes exhibit defects in phagocytosis caused by knockout of *RAGE*; for example, in HEK293T cells, the level to phagocytose apoptotic cells was decreased by 67% compared to that in wild-type cells when the phagocytosis assay was performed with a 0.5 h incubation. While professional phagocytes, such as macrophages, are equipped with various phagocytic receptors [2], defects in phagocytosis have also been observed in *Rage*^-/-^ primary macrophages. RAGE-mediated phagocytosis is induced by histones and PS [15, 22]. Since RAGE can recognize multiple molecules, other RAGE ligands, such as HMGB1 [38], may also serve as eat-me signals. Nevertheless, our results show that abrogation of histone-induced and RAGE-mediated phagocytosis causes a defect in the removal of apoptotic cells. For example, the number of HEK293T cells that phagocytose apoptotic cells was decreased by the removal of histones, when phagocytosis assays were performed with relatively short incubation times. Notably, however, when the assay was performed with a 1 h incubation, the phagocytosis of apoptotic cells by HEK293T cells was not inhibited by the removal of histones. Thus, histone-induced and RAGE-mediated phagocytosis is required for the rapid elimination of apoptotic cells. Given that PS is present in the plasma membrane, cell surface histones may be readily recognized by RAGE.

*Rage*^-/-^ mice exhibit defects in wound repair. During the wound healing process, neutrophils are removed from the wound site primarily via phagocytosis [33]. Clearance of these cells is required for resolution of inflammation; prolonged residence of the inflammatory cells in the wound site lead to the persistence of inflammation [39]. Thus, the delay in wound healing in *Rage*-KO mice may be attributable to the lack of RAGE-mediated phagocytosis. However, a previous report revealed that the activation of RAGE and the induction of proinflammatory signaling pathways can induce the proliferation and migration of cells [40, 41]. Thus, RAGE may be involved in wound repair in multiple ways. Nevertheless, the number of apoptotic neutrophils and histone-presenting neutrophils are increased at the wound site in *Rage*^-/-^ mice, suggesting that RAGE-mediated phagocytosis, including histone-induced processes, is required for the removal of apoptotic cells from the wound site. In addition to apoptotic cells, RAGE-mediated phagocytosis may be required for the removal of histone-containing macromolecules, such as NETs. The generation of apoptotic cells and NET formation are not merely limited to for the wound healing process. Thus, histone-induced, RAGE-mediated phagocytosis, and phagocytosis are involved in various processes to maintain healthy states in vertebrates.

## Materials and methods

### Cells

HEK293, HEK293T and Jurkat cells were obtained from American Type Culture Collection (ATCC). *RAGE*^-/-^ HEK293T cells were generated as described before [22]. HEK293T cells were cultured in Dulbecco’s modified Eagle’s medium (DMEM, Biological Industries) containing 10% (v/v) fetal calf serum (FCS) (Biological Industries). FCS was inactivated by incubation at 56°C for 45 min before addition to media. Jurkat cells were cultured in Roswell Park Memorial Institute (RPMI) 1640 (Biological Industries) containing 10% (v/v) FCS. Jurkat cells stably expressing GFP were cultured in RPMI 1640 containing 10% (v/v) FCS and 5 μg/ml blasticidin (InvivoGen). Mouse thymocytes were cultured in RPMI1640 medium containing 10% (v/v) FCS and 1% (v/v) penicillin-streptomycin solution (Beyotime). Primary lung epithelial cells were cultured in DMEM/F-12 medium containing 10% (v/v) FCS and 1% (v/v) penicillin-streptomycin solution, peritoneal macrophages were cultured in RPMI 1640 containing 10% (v/v) FCS and 1% (v/v) penicillin-streptomycin solution. All cells were maintained at 37°C in a humidified atmosphere with 5% CO_2_.

### Mice

*Ager*^+/-^ C57BL/6 mice purchased from Saiye Biology were bred at 6 to 10 weeks of age. The mice were housed in the specific pathogen free facility at the Experimental Animal Center of Jiangnan University. The housing conditions are as follows: the light time is 8:00-20:00, the temperature is 18-22°C and the relative humidity is 40-70%. Animal care and handling procedures comply with the National Institutes of Health Guidelines for the Care and Use of Laboratory Animals and are approved by the Ethics Committee of the Laboratory Animal Center of Jiangnan University. Mouse genotypes were determined by PCR using primers Ager-F1, Ager-R1, and Ager-R2 (Supplementary Table 1).

### Plasmids

All the oligo DNAs and plasmids used in this study are listed in Supplementary Table 1 and 2, respectively. For the CRISPR-Cas9 system used to knockout target genes, guide RNA sequences were designed using the E-CRISP website (RRID:SCR‒019088), and the corresponding DNA fragments were ligated into the *Bpi*I-digested vector pX330EGFP-hU6-gRNA-hSpCas9 [42]. The RAGE DNA fragments were amplified from cDNA derived from HEK293T cells and cloned into pME-Hyg-3FLAG [43]. pLIB2-mEGFP-BSD was used to make Jurkat cells stably expressing GFP. The GFP (mEGFP) was digested out of pME-mEGFP [22].

### Transfection

For transient transfection, cells were grown to 60% confluence in 6 cm plates and transfected with plasmid DNA (5 μg/well) using Lipofectamine 8000 (Beyotime) according to the manufacturer’s instructions.

Retrovirus-based transfection was performed as described previously [44] to construct Jurkat cells stably expressing GFP. HEK293T cells (10^6^) were transfected with 1 μg pGP, 1 μg pLC-VSVG, and 2 μg pLIB2-mEGFP-BSD using Lipofectamine 8000. After 36 h of incubation, the medium was filtered with a 0.22 μm filter and mixed with the same amount of DMEM supplemented with 16 μg/ml hexadimethrine bromide (Sigma‒Aldrich). The medium containing retrovirus was incubated with Jurkat cells overnight. After 5 days of culture, the cells expressing GFP were sorted using a cell sorter S3e (Bio‒Rad).

### Assays for RAGE and histones or DNA–histone binding

HEK293 cells were grown to approximately 70% confluence in 6 cm plates. The plasmids pME-RAGE^124-341^-His-FLAG or pME-RAGE^23-341^-His-FLAG were transfected into HEK293 cells. After two to three days of incubation, the medium was centrifuged at 3000 × *g* for 3 min at 4°C. The supernatant was loaded onto Ni NTA beads (Smart Life Sciences). His- and FLAG-tagged proteins were eluted with 400 mM imidazole. Subsequently, the anti-FLAG M2 affinity agarose gel (Sigma‒Aldrich) was added to the eluate and incubated for 3 h at 4°C with rotation. The agarose gel was washed with PBS (Sangon Biotech) three times and suspended in PBS. Two hundred microlitres of the agarose gel suspension was mixed with 1 mg/ml purified histone complex (Sangon Biotech) or 10 µg/µl recombinant histone H3C1 (Sangon Biotech). The final concentration of histones in the mixture was 20, 50, or 100 µg/µl, and the final concentration of recombinant H3C1 in the mixture was 1 µg/µl. The mixture was incubated at 4°C for 0.5 h. After washing three times with PBS, the agarose gel was suspended in 40 µl of PBS, 10 µl of 5 × SDS‒ PAGE loading buffer (250 mM Tris-HCl pH 6.8, 10% SDS, 0.5% bromophenol blue, 50% glycerol, and 5% ß-mercaptoethanol) was added, and the mixture was subjected to western blotting. To prepare the DNA-histone complex and DNA-histone H3.1, 1 nmol of a single-stranded 22-nt DNA (GTGCCAGATCGGGGTTCAATTC) fragment was incubated in 100 µl of 100 µg/µl histone or 100 µl of 1 µg/µl recombinant histone H3C1.

### Induction of apoptosis in Jurkat cells and thymocytes

Jurkat cells expressing GFP were resuspended in RPMI 1640 containing 10% FCS at a density of 10^6^/ml. The cells were treated with 1 μM staurosporine (Beyotime) for 12 h to induce apoptosis.

Thymus were harvested from 3 to 4 week-old wild-type C57BL/6 mice and chopped to produce a single-cell suspension. Then, the PKH67 Green Fluorescent cytomembrane Linker Kit (Solarbio) was used to label mouse thymocytes at 4°C for 20 min. To induce apoptosis, mouse thymocytes were re-suspended in RPMI1640 medium containing 10% FCS, 1% (v/v) penicillin-streptomycin solution supplemented with different concentrations of dexamethasone at 10^6^ cells/ml at 37°C for 12 h, and their viability was measured by Cell Counting Kit-8 (Sigma‒Aldrich) and the IC50 was calculated as described before [45]. And finally use 2.5 µM dexamethasone to induce mouse thymocytes apoptosis. Detection of apoptotic cells was performed with Apoptosis Detection Kit (Dojindo).

### Western blot analysis

Cells cultured in 6 cm plates were suspended in 500 μl RIPA buffer (Beyotime) supplemented with 5 µl of EDTA-Free proteinase inhibitor cocktails (MCE) and incubated at 4°C for 30 min. Protein concentrations in the supernatants were determined using BCA kit (Beyotime). 10 µg of each supernatant was mixed with 5 × SDS-PAGE loading buffer. After incubation at 100°C for 5 minutes, samples were subjected to 12% SDS-PAGE and transferred to PVDF membranes (Bio‒Rad). The membranes were blocked in 5% milk (Sangon Biotech) in TBST buffer (10 mM Tris-HCl, pH 7.5, 150 mM NaCl, and 0.05% (v/v) Tween-20). The following primary antibodies were used: rabbit anti-RAGE (Abcam, 1: 2000); rabbit anti-Caspase-3 (CST, 1:1000); rabbit anti-Cleaved caspase-3 (CST, 1:1000); mouse anti-actin (TransGen Biotech, 1:3000); mouse anti-Flag (TransGen Biotech, 1:5000) and rabbit anti-Histone 3 (Abcam, 1: 1000). The appropriate antibodies diluted in primary antibody dilution buffer (Beyotime) overnight at 4°C. After washing with TBST three times, the membranes were incubated with HRP-conjugated secondary antibodies diluted in 5% milk in TBST buffer at room temperature for 1 h and washed three times in TBST buffer. Primary antibodies were detected using the secondary anti-mouse IgG HRP-linked (TransGen Biotech, 1:5000), or anti-rabbit IgG HRP-linked (TransGen Biotech, 1:5000). Signals were detected with ECL Substrate (Bio‒Rad). Images were captured using a Tanon 5200 Automatic Chemiluminescence Image Analysis System.

### Immunofluorescence

Apoptotic Jurakt cells with GFP were induced as described above. Cells were washed with PBS twice and blocked with 1% bovine serum albumin in PBS for 30 min. The cells were incubated with anti-histone H3 antibody (1: 400) at 4°C for 1 h. Cells were washed with PBS three times, and were incubated with Alexa Fluor 555 donkey anti-rabbit IgG (InvivoGen, 1:1000) at 4°C for 30 min. After wash with PBS twice, the cells were observed under the microscope.

### Isolation and extraction of primary lung epithelial cells

For the purification and culture of primary lung epithelial cells, the lungs were perfused with buffered saline through the right ventricle and carefully resected. Two millilitres of dispase (Beyotime) were dripped into the lungs, followed by the addition of 1% low-melting agarose (Beyotime) and culture at 37°C for 1 h. After incubation, the lungs were minced and mechanically disrupted by passage through 100 and 70 µm nylon strainers. Single-cell suspensions were washed with PBS containing 1% FBS and 100 units of DNase I (Beyotime) and treated with red blood lysis buffer (Solarbio) at 4°C for 15 min. Then, the cell suspensions were centrifuged at 1000 × *g* for 5 min, washed twice with PBS, and treated with APC-conjugated anti-mouse CD45 antibody (BioLegend) and APC/Cy7-conjugated anti-mouse EPCAM antibody (BioLegend). CD45-negative and EPCAM-positive cells were collected via BD FACS Aria III, and the collected cells were resuspended in DMEM/F-12 medium supplemented with 10% (v/v) FCS and 1% (v/v) penicillin‒streptomycin solution. A total of 2 × 10^5^ cells were cultured in 24-well plates at 37°C.

### Isolation and preparation of mouse peritoneal macrophages

The cells were collected from mice peritoneal lavage after 8 to 10 weeks of age as described previously [46]. Macrophages (2.5 × 10^5^) were cultured in 24-well plate in RPMI 1640 medium supplemented with 10% (v/v) FCS and 1% (v/v) penicillin‒streptomycin solution. Cells were cultured in 24-well plates at 37°C, and after 1 h, the cells were washed with the medium to remove non-adherent cells.

### Treatments of apoptotic cells with enzymes, histones, and antibodies

To treat the apoptotic cells with DNase I or proteinase, 2 × 10^7^ apoptotic cells were incubated with 3 U of DNase I or 2 U of proteinase (Sigma-Aldrich) in PBS at 37°C for 30 min. Apoptotic cells were washed twice with PBS. Histone (final concentration, 5 µg/µl) or recombinant histone H3C1(final concentration, 5 µg/µl) were added to apoptotic cells and incubated at 4°C for 30 min to perform competitive inhibition assays. For treatment with antibodies, 2 × 10^7^ apoptotic cells were incubated with rabbit anti-histone H3 antibody (1: 400) or rabbit IgG monoclonal antibody-isotype control (Abcam, 1: 400) in PBS at 4°C for 30 min and washed twice with PBS.

### Phagocytosis assays

Phagocytosis assays were performed as described previously [22]. For the microscopy-based phagocytosis assay, round glass cover slips were placed on the bottom of 24-well plates. The cells (1.5 × 10^5^/well) were seeded in the plate and allowed to grow to a density of 2–3 × 10^5^ cells/well in 500 µl of media supplemented with 0.025 µl of LysoTracker (Beyotime). Apoptotic Jurkat cells were added to 10^6^ apoptotic cells/4–6 × 10^5^ HEK293T cells/ml media. The cells were incubated at 37°C for 30, 60, or 120 min. Then, the cells were placed on ice and washed twice with PBS, and the intracellular spores or beads were analyzed under a fluorescence microscope. Spores observed in LysoTracker-positive compartments were defined as internalized particles. At least a total of 300 cultured cells were analyzed.

A flow cytometry-based phagocytosis assay was performed as described before [46]. HEK293T cells were incubated with apoptotic Jurkat cells as described above. Primary lung epithelial cells and macrophages were cultured in 24-well plates as described previously. A total of 10^6^ apoptotic thymocytes were added to each well and incubated at 37°C for 0.5 or 1 h. The cells were washed with cold PBS to remove uninternalized cells. The cells were treated with trypsin and washed twice with PBS. The samples were analyzed via BD FACS Aria III. The data were analyzed via FlowJo X (BD).

For RAGE rescue experiments, wild-type cells were transfected with the pME-His empty plasmid, and *RAGE*^-/-^ HEK293T cells were transfected with the pME-His empty plasmid or the pME-RAGE-His plasmid.

### Microscopy

Microscopy images were obtained using a Nikon C2 Eclipse Ti-E inverted microscope with a DS-Ri camera equipped with NIS-Element AR software, and quantification of fluorescence intensity was performed using ImageJ software.

### Flow cytometry

Lung epithelial cells were sort by BD FACS Aria III, mice skin wound samples were analyzed by Beckman CytoFLEX S, and other samples were analyzed by Accurf C6 (BD). The data was analyzed using FlowJo V10.

### Generation of skin wounds in mice and analysis of wound tissues

Full-thickness punch wounds were made on the backs of the mice as described previously [47]. The mice were given analgesia and general anaesthesia, the back skin was shaved, the skin was lifted, and a mouse punch with a 3 mm aperture was made on the backs of the mice. Back skin samples of the same size were collected from the mice three days after wounding and incubated overnight with DMEM containing 1% (v/v) penicillin‒streptomycin solution and 200 µg/ml dispase II at 4°C. Small patches of skin were further digested with 1.5 mg/ml IV collagenase (Solarbio) and 10 U DNase I in RPMI 1640 medium supplemented with 2% FCS (v/v) and 1% (v/v) penicillin‒streptomycin solution. The suspension was subsequently resuspended in fresh digestion buffer and incubated at 37°C for 90 min. After incubation, the cell suspension was filtered with 70 µm nylon strainers to remove debris and clots. Neutrophils were sorted via PerCP-conjugated anti-mouse CD45 antibody, FITC-conjugated anti-mouse CD11b antibody (BioLegend, 101206), and APC-conjugated anti-mouse Ly-6G antibody (BioLegend). An annexin V-PE/7-AAD Apoptosis Detection Kit (Vazyme Biotech) was used to detect whole apoptotic cells, and an anti-histone 3 antibody was used to analyze cell surface histones.

### Statistical analysis

Each experiment was performed with at least three independent samples. The quantitative data were presented as mean ± Standard Error (SEM). Differences between the analyzed samples were considered significant at *P* < 0.05. Statistical significance was determined with two-tailed unpaired Student’s t test calculated with GraphPad Prism 8.4.3 software.

### Data availability

The data supporting the findings of this study are available within the Article and its Supplementary Information. Further relevant data are available from corresponding authors upon reasonable request.

## Acknowledgements

This work was supported by NSFC grants to H.N. (32071467) and X.Z. (32101031), and Open Foundation of Key Laboratory of Carbohydrate Chemistry and Biotechnology from Ministry of Education, China (KLCCB-KF202303).

## Author contributions

Conceptualization, H.N. and Y.L.; methodology, Y.L., X.Z., Y.Y., Y-S.L., and H.N.; validation, Y.L., Y.Y., C.D., Y-S.L., and H.N.; formal analysis, Y.L., X.Z., Y-S.L., and H.N.; investigation, Y.L., Y.Y., C.D., and Y-S.L.; writing—original draft preparation, Y.Y., X.Z. and H.N.; writing—review and editing, H.N., Y-S.L., and Z.L.; supervision, X.Z. and H.N.; funding acquisition, X.Z., Z.L. and H.N. All authors have read and agreed to the published version of the manuscript.

